# Toward Understanding the Role of Visual Hierarchy in Flicker-induced Time Dilation: A Preregistered Study

**DOI:** 10.1101/2025.11.10.687470

**Authors:** Amirmahmoud Houshmand Chatroudi, Yuko Yotsumoto

## Abstract

When a temporal interval is marked by flickering stimuli, its subjective duration is overestimated. The cortical origin of this temporal illusion, termed flicker-induced time dilation (FITD), is not yet well understood. In this pre-registered study, we aimed to explore whether the higher-level regions of ventral visual stream contribute to FITD. By using the semantic wavelet-induced frequency tagging (SWIFT) technique, four experimental conditions were created. Luminance and semantic flickers were created to selectively target the lower- and higher-level regions of the visual hierarchy, respectively. A combined luminance and semantic flicker condition was additionally created to probe the effect of simultaneous entrainment of lower- and higher-level visual regions on FITD. Lastly, to account for the single-pixel modulations and ripple-like motions induced by SWIFT, a control scramble condition was included. The phase-clustering analysis (N = 20) confirmed that the control scramble condition did not result in neural entrainment, whereas the flicker conditions showed entrainment whose magnitude and topography were consistent with their respective experimental manipulations. Nonetheless, the behavioral results indicated that the time dilation effect was not modulated by the flickering conditions neither was its magnitude correlated with the size of entrained oscillations. Moreover, the SWIFT scrambles led to a time dilation effect which was comparable to the flickering conditions. We discussed that while our results are not in agreement with neural entrainment account of FITD, they are not in agreement with the *processing principles* definition of salience either. Based on the putative shared cortical sources of SWIFT scrambles and luminance-modulated flickers, we conjectured that FITD, and motion-induced time dilation may rely on the same mechanism based on activation of motion sensitive regions. Finally, our results indicated that, at least at the presence of steadily activated lower-level visual and/or motion sensitive areas, periodic activation of ventral stream does not contribute to FITD.

## Introduction

Time is an abstraction inferred from sensory objects. There is no biological receptor that reacts to time and there is no sensory organ perceiving it. Sensory cortices in olfactory [1,2], visual [3–6], and auditory [7–9] systems encode time, and time perception depends on non-temporal sensory characteristics of objects (see [10] for a review). Temporal estimation *dilates* with an increase in the number of events presented within an interval and with the regularity with which these events are organized [11]. In vision (both for stationary flickers and moving objects [11–13]) and auditory modalities [14–16], it has been shown that regularly spaced stimuli expand the subjective time greater than anisochronous, accelerating and decelerating ones (but also see [17,18]). Thus, regularity is a salient property of sensory organization, and coincidentally, regularly spaced sensory stimuli are capable of entraining neural oscillations at their driving frequency [19–21]. The regularly spaced stimuli in vision, known as flickers, engender a range of temporal illusions marked by their ability in dilating the subjective time [5,22–25]. Nonetheless, the origin of flicker-induced time dilation (FITD) is a subject of debate. On one hand, salience-based accounts have shown that subjective rates of conscious perception of flickers correlate with the size of dilation [26]. On the other hand, neural entrainment accounts have established a relation between the amount of neural entrainment and the magnitude of the illusion [27]. The former is not easily falsifiable (due to variations in what one might consider salient; see Discussion), and support for the latter has been mixed [28,29].

Recently, we demonstrated that the residual excitatory cycles of an entrained 10-Hz oscillation, elicited by a rhythmic context, did not significantly impact the overestimation of subsequent flickers presented in-phase versus out-of-phase [29]. Thus, it was concluded that excitatory phases of entrainment may not play a major role in FITD. Consistent with this notion, the entrainment hypothesis falls short in explaining some other aspects of behavioral findings regarding FITD. For example, from the perspective of the neural entrainment hypothesis [26,27,30], it could be understood that the relation between FITD and temporal frequency should follow an inverse U-shape pattern, peaking at 8-12 Hz (due to the entrainment of alpha oscillations which have greater power and prevalence in EEG [20,30,31]). However, instead of an inverse U-shape, an ascending or descending pattern of FITD with frequency has been reported for sub-second [22] and supra-second ranges [26], respectively. The latter finding hints that the cortical oscillators which are engaged in FITD may be positioned in higher-level visual regions. This is because higher-level visual areas have poorer temporal resolution and lower temporal frequency sensitivity [32–35]. Therefore, when the duration is long enough (i.e., in the supra-second range), the slower frequencies can additionally entrain the higher-level visual areas [33]. The SSVEP topographic maps also align with this notion showing that at slower frequencies, distant cortical areas (such as frontal, parietal and temporal regions) coactivate with the occipital regions [20,36–42], yet as temporal frequency increases, the SSVEP sources get restricted to the primary visual areas [32,37–40,43]. Hence, it is conceivable that the prominent oscillators contributing to FITD are situated in higher-order visual areas, resulting in a pronounced time dilation at slower frequencies within the supra-second range. This may also explain why in the previous study [29], we failed to establish a relation between entrainment and FITD as we were focused on excitatory phases of entrained oscillation in primary visual areas.

To evaluate this hypothesis, we focused on the potential role of higher-level ventral visual regions in FITD. This investigation was informed by previous research on time distortions that qualitatively share similarities with FITD. Such studies, in explaining the time dilation effect induced by flickers, occasionally have resorted to ventral stream-based justifications. For example, van Wassenhove et al. [44] have attributed the time dilation of looming stimuli (expanding discs) to the threatening semantics such stimuli may carry in self-referential processes [45]. Similarly, Binetti et al. [46] have linked the opposite temporal estimation errors of accelerating and decelerating flickers to their gestalts (the spatio-temporal organization of flickers). Thus, to the degree the objecthood of flickers may matter [47], regularly spaced flickers may also additionally benefit from the contributions of the ventral stream (besides the implied prominent role of primary visual areas and the dorsal stream [48–51]). Thus, such semantic-based salience effect might be the missing link between the entrainment and salience theory of FITD, assuming that flickers, due to their regularity, are semantically categorized as salient percepts (see Discussion).

Supporting this claim, neuroimaging studies have reported that ventral visual regions contribute, albeit minimally, to the generation of SSVEPs elicited by flickers [41]. The ventral stream of the vision has classically been associated with the processing of visual properties such as high spatial frequencies, color, form, object recognition and consciousness [52,53] (see [54] for a review). Relatedly, the role of ventral stream in time perception has long been neglected. Nonetheless, in recent years ventral stream-based time dilation effects have started to surface (faces, [55], scenes [56], perceptual complexities, [57]) highlighting the role of perceptual content in time estimations [58]. Interestingly, a recent study has shown that the more memorable scenes tend to be judged as longer [59], hinting higher-order ventral areas such as inferotemporal cortex (IT) which selectively react to object categories [60] may play a more prominent role in time perception than what was envisaged before [61].

Thus, in this preregistered study, we aimed at exploring whether IT—implicated in the dilation of subjective time [59] and located downstream in the ventral pathway [60]—affects FITD. We reasoned if higher-level visual areas along this pathway play a major role, moving up the location of the entrainment from primary visual areas to the IT should amplify the dilation. In doing so, we exploited the newly introduced method of semantic wavelet-induced frequency-tagging (SWIFT [33]). By conserving low-level visual properties across the frames, SWIFT scrambles the local contours’ orientations so that the original image is shown in a periodic manner (at a base frequency). Thus, unlike classic luminance-based flickers, SWIFT has the selectivity and specificity to periodically activate higher-level ventral regions (in particular, IT) without entraining lower-level visual areas (i.e., lower-level visual areas are steadily activated [62]. Adopting this technique, the primary goal of this study was to assess whether semantic flickers, targeting higher-level regions of the ventral stream, can induce pronounced time dilations. Nonetheless, as a byproduct of the wavelet scrambling, the SWIFT sequence undergoes individual pixel modulations across frames. To control this effect, a condition was created that included SWIFT scrambling, but no semantic image was shown. Importantly, this condition is not anticipated to result in entrainment at any level of the visual hierarchy. Rather, owing to the normalized modulation of single pixels in this condition, a steady activation of lower-level visual areas is expected [62]. Moreover, due to the presence of random ripple-like motions [33], a non-entraining activation of motion-sensitive areas is also probable. In addition to the scramble condition, to compare semantic-based FITD with classic FITD, we created luminance-modulated flickers from SWIFT scrambles. Furthermore, by combining the SWIFT and classic luminance-based flickers (Hierarchical Frequency Tagging technique [63]), we sought to explore whether the coactivation of lower- and higher-level visual areas results in maximal FITDs. Lastly, by recording electroencephalography (EEG) we attempted to unravel whether phase clustering in SWIFT, luminance, or the combined flicker conditions display a clear correlation with the FITDs.

## Method

### Subjects

Twenty university students (mean age = 25.6, SD = 5.51, male = 17) participated in the experiment. The sample size was calculated based on a priori power analysis as it is detailed in the preregistration draft (osf.io/ec9qa). Moreover, data from one subject was removed from further analysis due to poor temporal sensitivity as it was set as an exclusion criterion in the preregistration plan. All subjects had normal or corrected-to-normal vision, with no medical history of seizure, or epilepsy. Subjects were paid for their participation, and written informed consent was obtained from them prior to the experiment. The experiment was conducted in accordance with the guidelines set by the ethics boards of the University of Tokyo (protocol #989, data collection period: 2024-08-05 to 2024-09-29).

### Apparatus

EEG data was collected using EGI Net Amps 400 amplifier (Geodesic EEG Systems, USA) with a standard 64 HydroCel GSN cap. The sampling rate was 1000 Hz, and the impedance was kept below 30 kΩ. The EEG data were online referenced to Cz and no online filter was applied to the data. The stimuli were created in MATLAB (MathWorks Inc, v2024a) environment using Psychtoolbox-3 extension [64–66] and were presented on a 23.6-inch VIEWPixx 3D monitor (VPixx Technologies, Inc., Canada) with refresh rate of 120 Hz. The monitor display was set to the scanning backlight mode which enables CRT-like behavior with precise timing and high inter-frame independence. Participants were seated in a reclining chair at a viewing distance of 70 cm from the monitor in a dark, soundproof room.

### Stimuli

Sixteen free-to-use images were found on Pexels website (https://www.pexels.com/) and were downloaded (four female faces, four male faces and eight houses). Our rational for using only house and face images was that the previous study using fMRI demonstrated that SWIFT can most effectively activate higher-level (ventral) visual regions using houses and faces, and not objects [62]. Next, the images were transformed to grayscale (0 to 1 range), resized to 300 × 300 pixels (6.7°) and were equalized in contrast using histogram equalization function (histeq.m) in MATLAB. Subsequently, the images were equalized for average luminance. Next, SWIFT cycles were created using SWIFT function [33]. The number of harmonics used was three, and the level decomposition was nine. SWIFT is a technique which, by decomposing an image using wavelets and rotating the contours along an isoenergetic sphere, can create scramble versions of the image while conserving the local energy and spatial frequency distributions (for a complete description of the method, see [33,62]). The rotation of contours occurs at a fundamental frequency (f0) and its harmonics, which ensures the original image contours are periodically realigned at f0. Since the highest phase clustering using SWIFT was achieved at 2 Hz [33], to achieve the maximal phase clustering in this study, we set the f0 to 2 Hz (consequently, the frequency of all flickering conditions was 2 Hz). For each image, multiple SWIFT cycles were created (the length of each SWIFT cycle was 0.5 seconds). Subsequently, the relative images similarity (RIS) index for the frames in each cycle was calculated. The RIS for each frame is calculated by averaging the absolute distances of the wavelet coefficients in that frame from the coefficients of the original image (for a detailed description, see [62]). This value is then inverted and scaled so that the most similar frame has a value of 100%, while the most dissimilar frame has a value of 0%. By using RIS index for each image, eight SWIFT cycles which showed a clear U shape in their similarity pattern (i.e., cycles with no sudden realignment of harmonics in the middle of SWIFT period leading to momentarily increase in RIS) were selected (e.g., Fig 1a). Using the SWIFT sequences, five conditions were created as follows:

**Fig 1.**
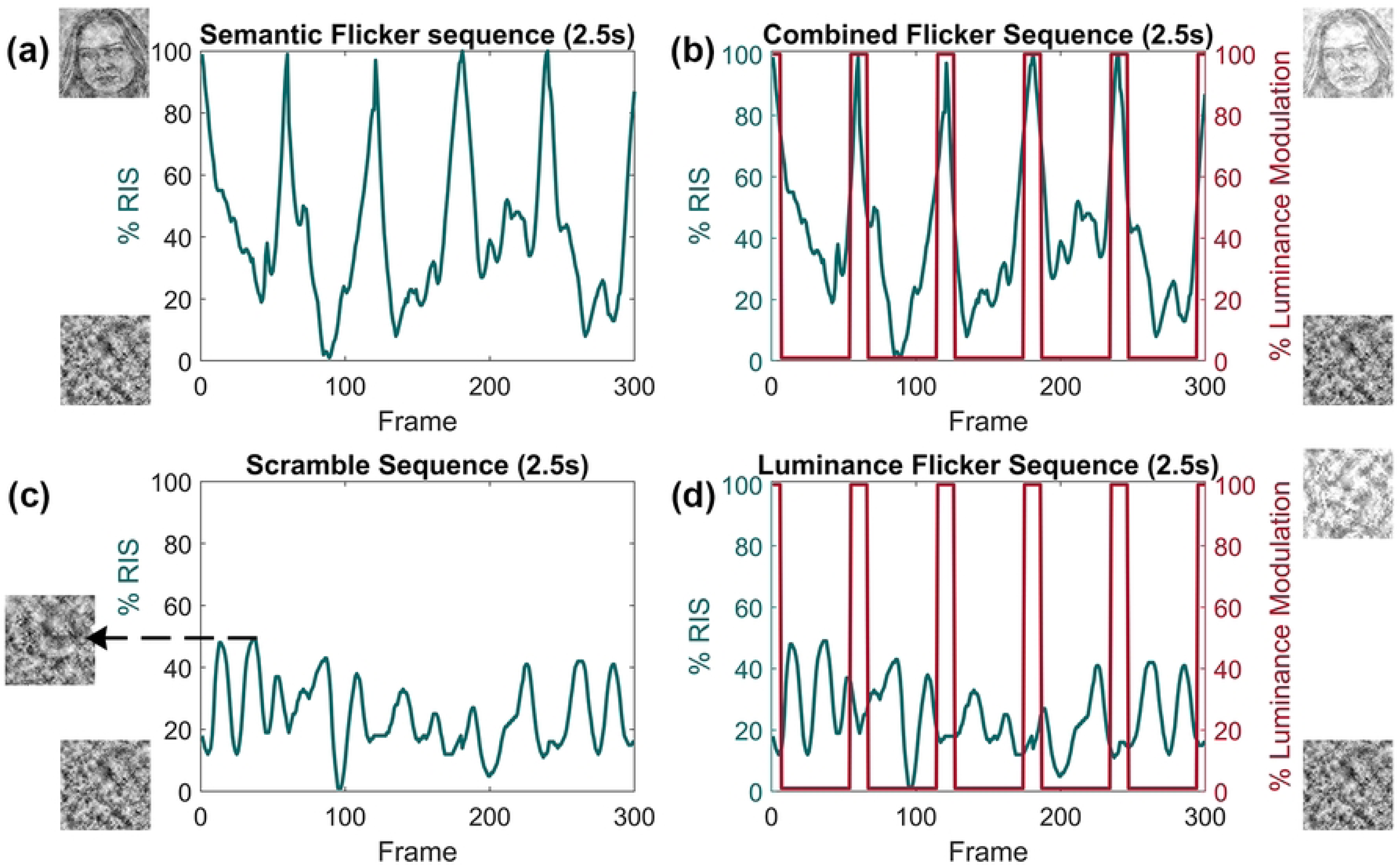
Examples of SWIFT sequences used in experimental conditions. (a) In the semantic flicker condition, on each trial five random cycles (out of eight) were concatenated. RIS refers to the relative image similarity index which indicates the similarity of each sequence’s frame with the original image. 0% RIS represents total scramble and 100% RIS indicates a recognizable image. During a SWIFT sequence, the original image becomes recognizable on a few frames before it fades into scrambling again. This gives SWIFT cycles a typical u-shape pattern [62] which periodically activates higher-level ventral visual regions. (b) In the combined flicker condition, a luminance modulation was applied on top of the semantic flickers. Importantly, this luminance modulation occurred in-phase with the RIS index. This means on the few frames where image was recognizable, the luminance was also increased. This condition was intended to periodically activate both lower- and higher-level visual areas (Hierarchical Frequency Tagging [63]). (c) Scramble sequences for each image were created by taking the most dissimilar frame from semantic sequences in (a) and feeding it again to the SWIFT function. On each trial, five random scrambled cycles (out of eight) were concatenated. As the RIS index indicates, the scrambles’ similarity fluctuated but never became a recognizable image. (d) In the luminance flicker condition, a luminance modulation was applied on top of scramble sequences. This condition was intended to periodically activate low-level visual areas. The frequency of modulation in all conditions was 2 Hz.

#### Semantic flicker condition

on each semantic flicker trial, for each image, five out of eight SWIFT cycles were chosen uniformly at random and were concatenated to make a 2.5-second-long sequence. Next, each sequence underwent a normalization procedure. First, the contrast was equalized across frames using contrast-limited adaptive histogram equalization (adapthisteq.m) function in MATLAB. Next, pixels across frames were converted into the frequency domain using the Fourier transform. Following that, the aggregated individual pixel frequency spectra was equalized in amplitude and was cut off at the half distance of harmonics after the last harmonic of f0 (the individual pixel frequency spectra’s cut-off in this study for 2.5-second-long sequences at f0 of 2 Hz was 7.2 Hz, Fig 3a). This normalization ensured that the individual pixel modulations did not have an increased power at a particular frequency and rather all pixels were modulated in a white noise manner (i.e., having identical spectra amplitude). Lastly, each frame was equalized in global luminance to 0.5 gray scale (Fig 3b). This normalization procedure ensured that in addition to the spatial frequencies (due to isoenergetic wavelet rotation), all low-level visual features (contrast, individual pixel modulations, global luminance modulations) were the same across the frames. Yet the original image was periodically presented at 2 Hz (Fig 1a; see supplementary materials Movie S1, for an exemplary semantic flicker sequence).

#### Scramble condition

for each image, from the already-created eight SWIFT cycles, the frame that had the lowest similarity to the original image was chosen (based on RIS index). Next, this frame was fed to the SWIFT function to create eight new SWIFT cycles. These cycles did not have the original image embedded in them; thus, they were just a scrambled version of the semantic flickers. On each trial corresponding to the scramble condition, five out of the eight scramble cycles were chosen uniformly at random and were concatenated to make 2.5-second-long sequences (Fig 1c; Movie S2). Next, these sequences underwent the same normalization procedure as explained for the semantic flickers (Fig 3a & b).

#### Luminance flickers

on each luminance flicker trial, first a 2.5-second-long scramble sequence was created as explained above. Next, the global luminance of these sequences was modulated in a square wave manner (12 frames on and 48 frames off, modulation depth = +0.25 grayscale; Fig 1d, Fig 3a & b; Movie S3).

#### Combined flickers

on each combined flicker trial, first a 2.5-second-long semantic flicker sequence was created as explained above. Next, the global luminance of these sequences was modulated in a square wave manner (12 frames on and 48 frames off, modulation depth = +0.25 grayscale). Importantly, the luminance and the semantic modulations were in-phase, meaning the on-period of increased luminance always accompanied the few frames where the original image was presented (Fig 1b, Fig 3a & b; Movie S4).

#### Static condition

on each trial corresponding to the static condition, for each image, from the randomly selected five SWIFT cycles, the frame which had the highest similarity to the original image was chosen (based on RIS index). This image was statically presented for the duration of 2.5 seconds (Fig 2).

**Fig 2.**
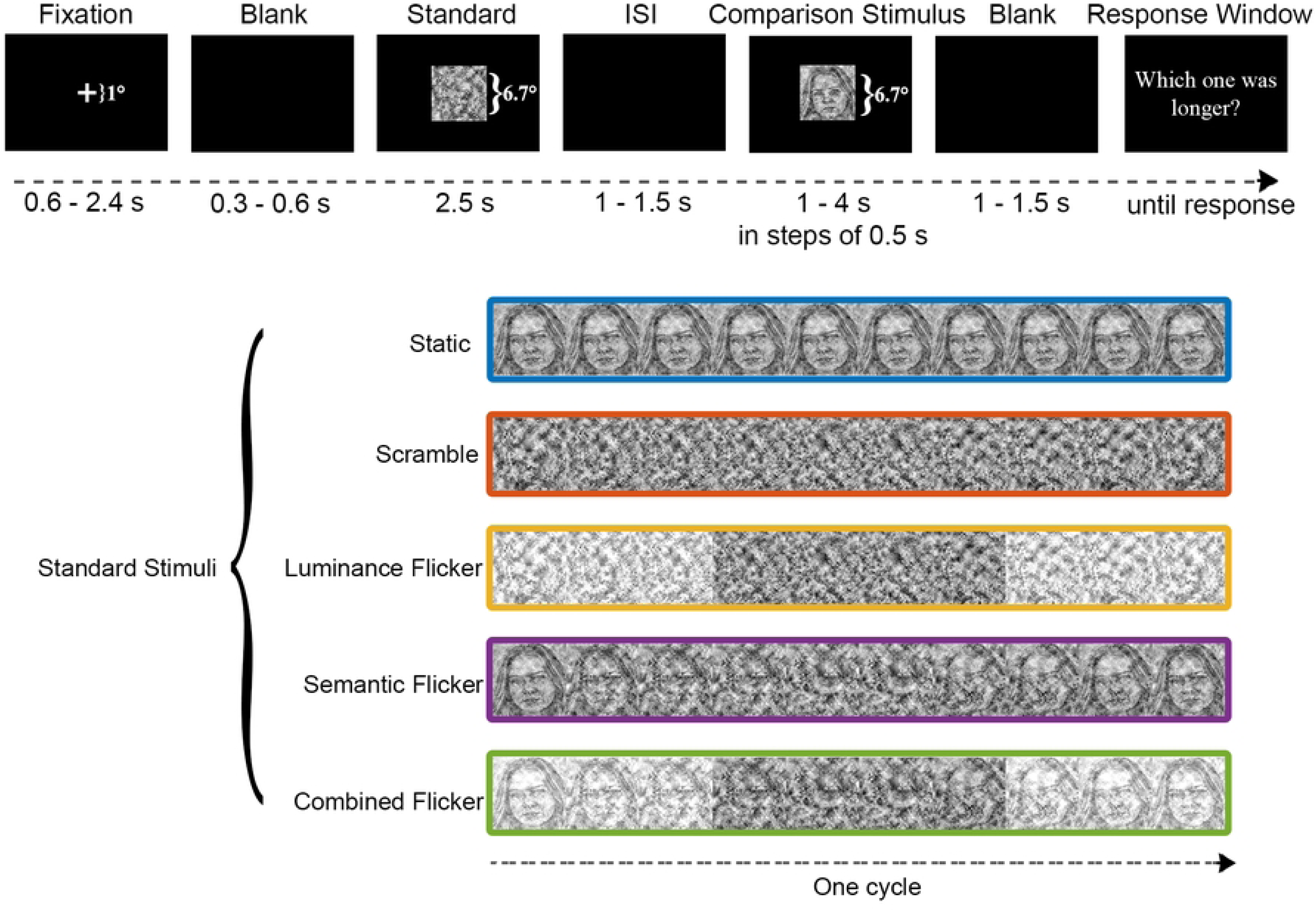
Duration Comparison task. On each trial, following the fixation, either a standard or a comparison stimulus was shown. Subjects were asked to choose the stimuli that lasted longer. The comparison stimulus was always a static image which could have a random duration from 1 to 4 s in steps of 0.5 s. The standard stimuli had a fixed duration of 2.5 s. The standard stimulus could have five conditions which are shown on the bottom of the panel. The order of standard and comparison stimuli, durations, and standard stimuli’s conditions were randomized. On each trial the static and comparison stimuli were derived from same image to control for low-level visual and high-level semantic confounders.

**Fig 3.**
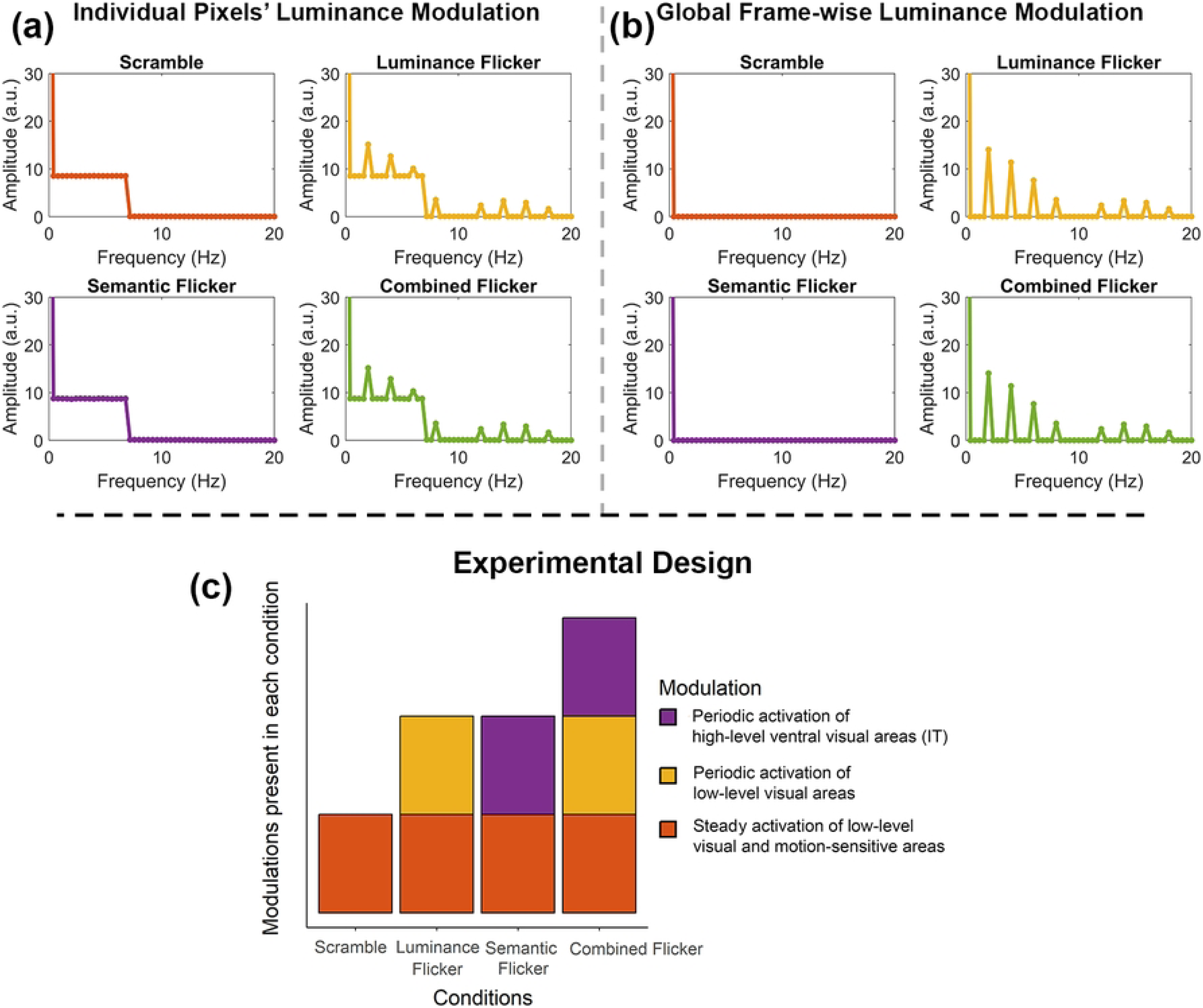
Experimental conditions’ frequency modulation spectra and the experiment’s design. (a) In semantic and scramble conditions, the SWIFT normalization ensured that the individual pixels were modulated equally across a definite frequency spectrum (cut-off frequency in this experiment was 7.2 Hz, see text) with no peaks at any single frequencies. (b) In luminance and combined conditions, the luminance of frames was periodically modulated (Fig 1b & d) at 2 Hz. However, in semantic and scramble conditions, the mean luminance across frames was equalized. Thus, no frame-wise modulation is seen for these conditions. (c) Due to constant rotation of the image contours by SWIFT function and consequently, the normalized single-pixel luminance modulations, the lower-level visual areas are steadily activated [62]. Moreover, the rotation of the image contours induces ripple-like motion [33], thus putatively activating motion-sensitive areas. This steady activation is shared across all experimental conditions. In the luminance flicker condition, due to modulation of luminance, the low-level visual areas are periodically activated. In the semantic flicker condition, due to realignment of image contours, the original image periodically becomes recognizable. Thus, the higher-level ventral regions (in particular, IT) are consequently periodically activated [62]. In the combined flicker condition, due to the simultaneous modulation of luminance and semantic information, both low- and high-level visual areas are periodically activated [63]. In this experimental design, we probed whether the magnitude of FITD is dependent on periodic activation of higher- vs lower visual areas and whether it will show a notable correlation with indices of neural entrainment (phase clustering) in different parts of the visual hierarchy.

### Task

The changes in perceived subjective durations across conditions were measured using a duration comparison task (Fig 2). Each trial began with a fixation period with random duration ranging from 0.6 to 2.4 seconds, followed by a random blank period of 0.3 to 0.6 seconds. Subsequently, a standard stimulus and a comparison stimulus were sequentially presented with a random inter-stimulus interval (ISI) ranging from 1 to 1.5 seconds. After a random blank period of 1 to 1.5 seconds, subjects were asked to choose the stimuli that lasted longer in a 2-interval forced choice design (2IFC). The duration of the comparison stimulus was 1, 1.5, 2, 2.5, 3, 3.5, and 4 seconds. The order of the standard and comparison stimuli was counterbalanced across conditions, images, and comparison durations. The duration of the standard stimulus was 2.5 seconds. The standard stimulus had five conditions as explained above (Fig 2). The comparison stimuli were always static, and they were created in an identical manner to the standard static condition. It is important to note that on each trial, the same image was used both for the standard and comparison stimuli (i.e., the same image was used as a static comparison stimulus and was also fed to create semantic, scramble, luminance, combined, or static standard stimuli). This was intended to control low-level visual and high-level semantic confounders between standard and comparison stimuli. The stimuli were presented on a black background. The size of the fixation cross was 1°.

### Procedure

Subjects were asked to focus on the stimuli presented at the center of the screen, refraining from any muscle or eye movements during the stimuli’s presentation time. Importantly, they were asked not to count the stimuli’s duration and choose the interval that lasted longer. A total of 16 (images)×7 (durations)×5 (conditions) trials were presented. Therefore, for each condition, and each duration, each image was repeated only once. The total of randomized 560 trials were segmented into seven blocks with six mandatory rests in between (each 5 to 8 minutes). The experiment lasted three hours.

### Analysis

#### EEG data

Preprocessing: EEG data were first low-pass filtered at 30 Hz and subsequently, high-pass filtered at 0.5 Hz using EEGLAB (version 2024.2 [67]). Next, 4-second-long epochs (0.5-second baseline) were created and were baseline corrected. Next, since the peripheral electrodes in our setup were of no use, they were removed from the EEG data (E1, E17, E23, E29, E32, E43, E47, E55, E61, E62, E63, and E64 channels were discarded). Following that, the EEG data was visually inspected, and noisy channels in each epoch were interpolated. Epochs with excessive noise were removed. This resulted in removal of 4% of trials which were not significantly different across conditions (χ (4) ^2^ = 0.93, *p* = 0.91). Moreover, an independent component analysis was done to remove eye and muscle artifacts. Lastly, the data were average referenced.

#### ITPC

Time-frequency decomposition: to extract the phase of EEG data at each frequency, time point, and trial, EEG data were decomposed using complex Morlet wavelets at frequencies from 1 to 10 Hz in steps of 0.2 Hz using custom-written MATLAB scripts [68]. The length of the Morlet wavelet was set to eight seconds (-4 to 4 seconds centering on zero) and the number of wavelet cycles was set to the logarithmically spaced range of 6 to 8 to balance the time-frequency precision [68].

##### ITPC

for each frequency, the extracted phase values at each time point were vector-averaged across trials according to the following formula [68] (p. 244):

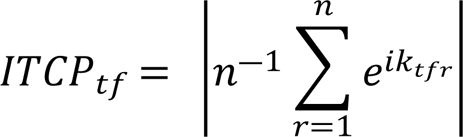

##### Hypothesis-driven analysis

ITPC analysis was restricted to a 1 to 2-second time window (relative to stimulus onset) by discarding the ITPC values at the beginning and end of the epochs [42]. We restricted our analysis solely to this time window to avoid transient phase clustering that occurs at the onset and/or offset of stimulus sequences. Moreover, the lower bound of this window was specifically chosen to be sufficiently far from the onset to allow a stable phase clustering to be detected (in accordance with previous studies [29,37,38,69]). Lastly, similar to previous studies using SWIFT [33,63,70], we limited the ITPC analysis to a broader visual area, containing occipital, parietal, and temporal regions by averaging ITPC values over E16, E20, E21, E22, E25, E26, E27, E28, E31, E33, E34, E35, E36, E37, E38, E39, E40, E41, E42, E45, E46, E48, E49, E50, E51, and Cz channels (Fig 4b). The ITPC values at 2 Hz were averaged over the selected channels and the time window regions of interest (ROI) for each subject. Next, these average values were probit transformed (i.e., normalized, as explained in the preregistration draft) and were subjected to a one-way repeated measures analysis of variance (rANOVA). Significance was followed by Bonferroni-corrected post hoc t-tests.

**Fig 4.**
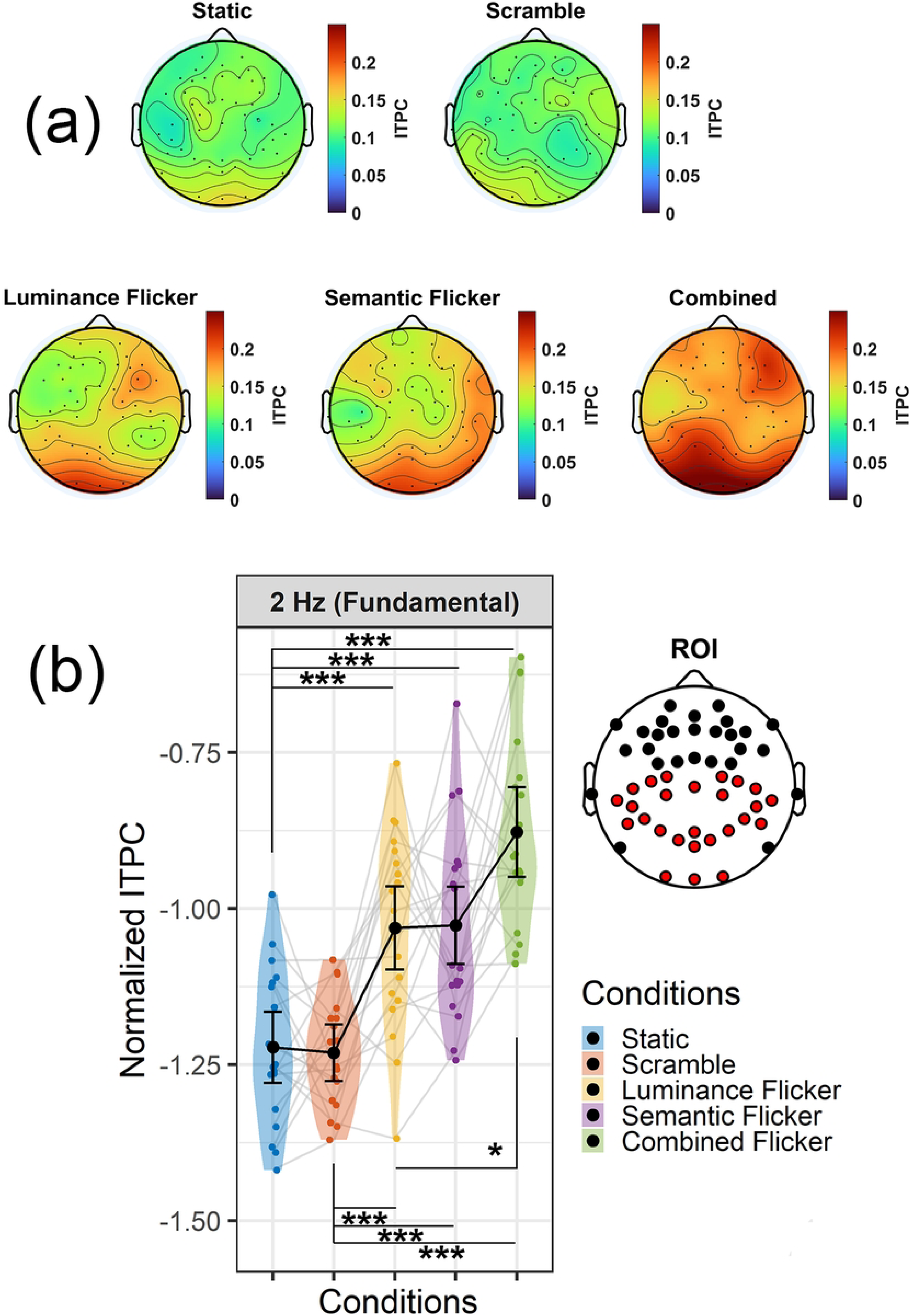
Topographic maps and hypothesis-driven analysis of ITPC data. (a) shows topographic maps of 2-Hz ITPC values. In static and scramble conditions, no notable phase clustering was observed. For luminance flickers, primary visual areas showed the major phase clustering. Semantic flickers resulted in a right lateralized phase clustering in occipitotemporal regions consistent with semantic processes of faces and houses (see text). The combined flicker condition resulted in a widespread phase clustering across the visual hierarchy. (b) Hypothesis-driven analysis of normalized 2-Hz ITPC values. The head plot shows the electrodes selected as the region of interest (ROI) by red color. The ITPCs in ROI electrodes were averaged over a window of 1 to 2 s relative to the sequence onset. There was no significant phase clustering difference between static and scramble condition, affirming that scrambles did not entrain 2-Hz neural oscillations. ITPC was significantly higher for flickering conditions, signaling entrainment had occurred. There was no significant ITPC difference between luminance and semantic flickers, implying comparable entrainment in lower- and higher-level visual areas. The combined condition resulted in stronger phase clustering than luminance flickers (it also showed a trend relative to semantic flickers, see text). Error bar indicates 95% confidence interval. * p<0.05, *** p<0.001.

##### Exploratory analysis

as explained in the preregistration draft, to explore whether the experimental task conditions containing 2 Hz flickers (semantic, luminance, and combined flicker conditions) had resulted in phase clustering other than that of interest (2 Hz), we performed time-frequency permutation statistical analysis. More importantly, this analysis was also intended to ensure that the scramble condition has not resulted in phase clustering at any frequency range. Thus, eliminating the possibility that any behavioral differences observed in this condition could be attributed to selective entrainment at frequencies other than that of 2 Hz. To this end, we performed the exploratory analysis over the 1 to 2-second time window (relative to onset sequence) and 1 to 10 Hz frequency range on the averaged ITPC values from the electrodes specified in the hypothesis-driven analysis. ITPC values in each experimental condition were separately contrasted against the static condition (i.e., the baseline condition). The significance of the observed contrast was then assessed using permutation-based testing. For this purpose, a null hypothesis distribution was generated by flipping the sign of the contrast for a random subset of subjects for 5000 times and storing the group-level average contrasts [68]. Next, to correct for the multiple comparisons, a pixel-based approach was adopted. In this conservative approach, each observed time frequency point is compared against the 95 percentiles of the maximum pixels of the null hypothesis distribution (p < 0.05, one-way) and the observed values higher than the specified percentile are marked as significant (for the detailed explanation of permutation-based statistical testing and pixel-based multiple comparison correction method refer to [68]).

#### Behavioral data

For each participant and for each condition a logistic function was fitted using Palamedes toolbox (v1.11.11 [71]). The lapse rate was set at 0.02 to account for the potential lapses of attention occurring due to the length of the experiment [72]. Moreover, the guess rate was set to zero. The point of subjective equality (PSE) and just noticeable difference (JND) were extracted and separately were subjected to rANOVA as detailed in the preregistration plan. Moreover, Bayse factors were obtained using one-way rANOVA in JASP [73].

Moreover, the relationship between PSE and 2-Hz ITPC values was analyzed using multiple linear regression in R with conditions set as one categorical factor (selecting the static condition as the baseline) and probit-transformed ITPC values as the other continuous independent variable. The purpose of this analysis was to find significant interactions between conditions and ITPC (i.e., evaluating whether the relation between PSE and ITPC is the function of conditions). A significant interaction was followed by post-hoc single regressions between ITPC and PSE, separately performed for each condition with false discovery rate [74] for multiple comparison corrections.

## Results

### EEG

Topographic maps of ITPC values at 2 Hz are shown in Fig 4a. As expected, the static and scramble conditions did not result in any phase clustering. The luminance condition showed a major phase clustering at the primary occipital region with some additional clustering at right frontal region, reflecting a prototypical luminance-based SSVEP profile [20,37–39]. The semantic condition resulted in phase clustering at occipital, right occipitotemporal, and right temporoparietal electrodes. This pattern is consistent with previous studies [63,70] using SWIFT, and the lateralization aligns with the preferential semantic processing of houses, and specially faces in the right occipitotemporal areas (see Discussion). Lastly, the combined flicker condition resulted in major phase clustering over a wide range of occipital, occipitotemporal, temporoparietal, and right frontal regions. This indicates that the combined condition periodically activated both low- and higher-level visual regions.

The hypothesis-driven ITPC analysis revealed that the magnitude of phase clustering at 2 Hz was not different between the static baseline and scramble condition (t (18) = 0.27, *p* = 1), affirming that scrambles did not result in phase clustering at this frequency (Fig 4b). Moreover, the 2-Hz phase clustering was highest for the flickering conditions relative to the static baseline condition (t (18) = 4.15, *p* = 0.006; t (18) = 5.83, *p* < 0.001; t (18) = 7.55, *p* < 0.006 for luminance, semantic, and combined flicker conditions, respectively). Similarly, the 2-Hz phase clustering was highest for the flickering conditions relative to the scramble condition (t (18) = 5.81, *p* < 0.001; t (18) = 5.64, *p* < 0.001; t (18) = 8.52, *p* < 0.006 for luminance, semantic, and combined flicker conditions, respectively). The amount of phase clustering was comparable between luminance and semantic flickers (t (18) = 0.09, *p* = 1), indicating the size of entrainment at lower and higher regions of the visual hierarchy was comparable (Fig 4b). Lastly, as expected, the phase clustering in the combined flicker condition was significantly higher than luminance flickers (t (18) = 3.47, *p* = 0.02), and showed a trend relative to semantic flickers (t (18) = 3.13, *p* = 0.058). Overall, these results confirmed that the scramble condition did not entrain 2-Hz oscillations relative to the static condition while a robust entrainment for flickering conditions relative to the static condition was observed. Moreover, the magnitude of the entrainment in the combined flicker conditions was greater than luminance-only and semantic-only flickers, reflecting the broader engagement of lower- and higher-level visual regions. Furthermore, in a follow-up analysis, we applied the same procedure to the harmonic frequencies and found qualitatively similar findings (see supplementary materials).

The exploratory ITPC analysis confirmed the results obtained from the hypothesis-driven approach. As it is shown in Fig 5, scramble condition did not result in any significant phase clustering between 1 to 10 Hz (the cut-off frequency of individual pixel’s modulation was 7.2 Hz, Fig 3a). As anticipated, a major phase clustering was observed for the luminance flicker condition at 2 Hz. A consistent phase clustering was also observed for the semantic flickers at 2 Hz albeit losing momentum towards the end of the presentation. Additionally, for semantic flickers, a significant phase clustering was also detected at 4 Hz (the first harmonic frequency). Combined flickers resulted in robust phase clustering at the modulation frequency (2 Hz) and its harmonics (4,6, and 8 Hz). The exploratory ITPC results thus aligned well with the hypothesis-driven findings further confirming that scramble condition did not entrain any neural oscillations while flicker conditions did. Moreover, the entrainment elicited by the combined condition was more robust, influencing a greater number of harmonic frequencies compared to the luminance-only and semantic-only conditions.

**Fig 5.**
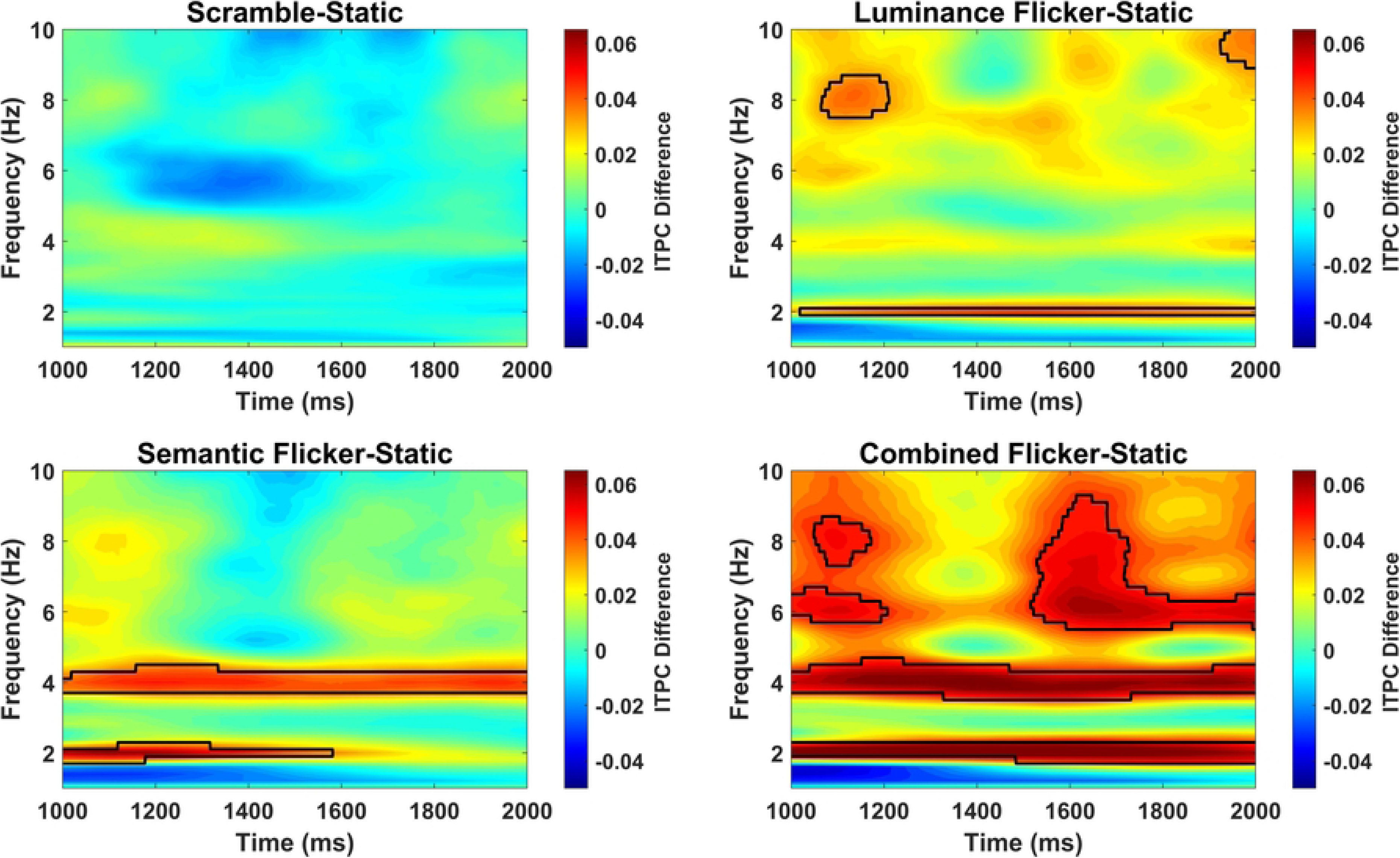
Time-frequency ITPC differences between experimental conditions and the static baseline condition. Clusters which were significant at p < 0.05, are demarcated by black lines. No significant ITPC cluster was detected in the scramble condition, indicating entrainment did not occur in this condition. In flickering conditions, a significant phase clustering was observed at 2 Hz, implying entrainment had occurred at these conditions. Additionally, in the semantic flicker condition, the first harmonic (4 Hz) and in the combined condition, first (4 Hz), second (6 Hz), and third (8 Hz) harmonics showed significant ITPC (for analysis of harmonic frequencies, see supplementary materials).

### Behavioral

The behavioral data (Fig 6a) indicated that the subjective duration of the flickering conditions was significantly overestimated relative to the static baseline condition (time dilation effect) as it is indicated by significantly higher PSEs for luminance (t (18) = 5.49, *p* < 0.001, BF01 = 0.001), semantic (t (18) = 5.44, *p* < 0.001, BF01 = 0.001) and combined conditions (t (18) = 6.78, *p* < 0.001, BF01 = 1.21e-4) relative to the static condition. Moreover, the size of time dilation was comparable across flickering conditions as the PSE was not significantly different between luminance and semantic (t (18) = 0.97, *p* = 1, BF01 = 2.77), luminance and combined (t (18) = 0.04, *p* = 1, BF01 = 4.2), and semantic and combined conditions (t (18) = 0.8, *p* = 1, BF01 = 3.17). Nevertheless, the scramble condition resulted in significant time dilation as the PSE in this condition was significantly higher than static baseline (t (18) = 5.66, *p* < 0.001, BF01 = 9.5e-4). Moreover, the size of dilation in this condition was comparable to luminance (t (18) = 1.05, *p* = 1, BF01 = 2.59), semantic (t (18) = 0.49, *p* = 1, BF01 = 3.77), and combined flicker conditions (t (18) = 1.15, *p* = 1, BF01 = 2.35).

**Fig 6.**
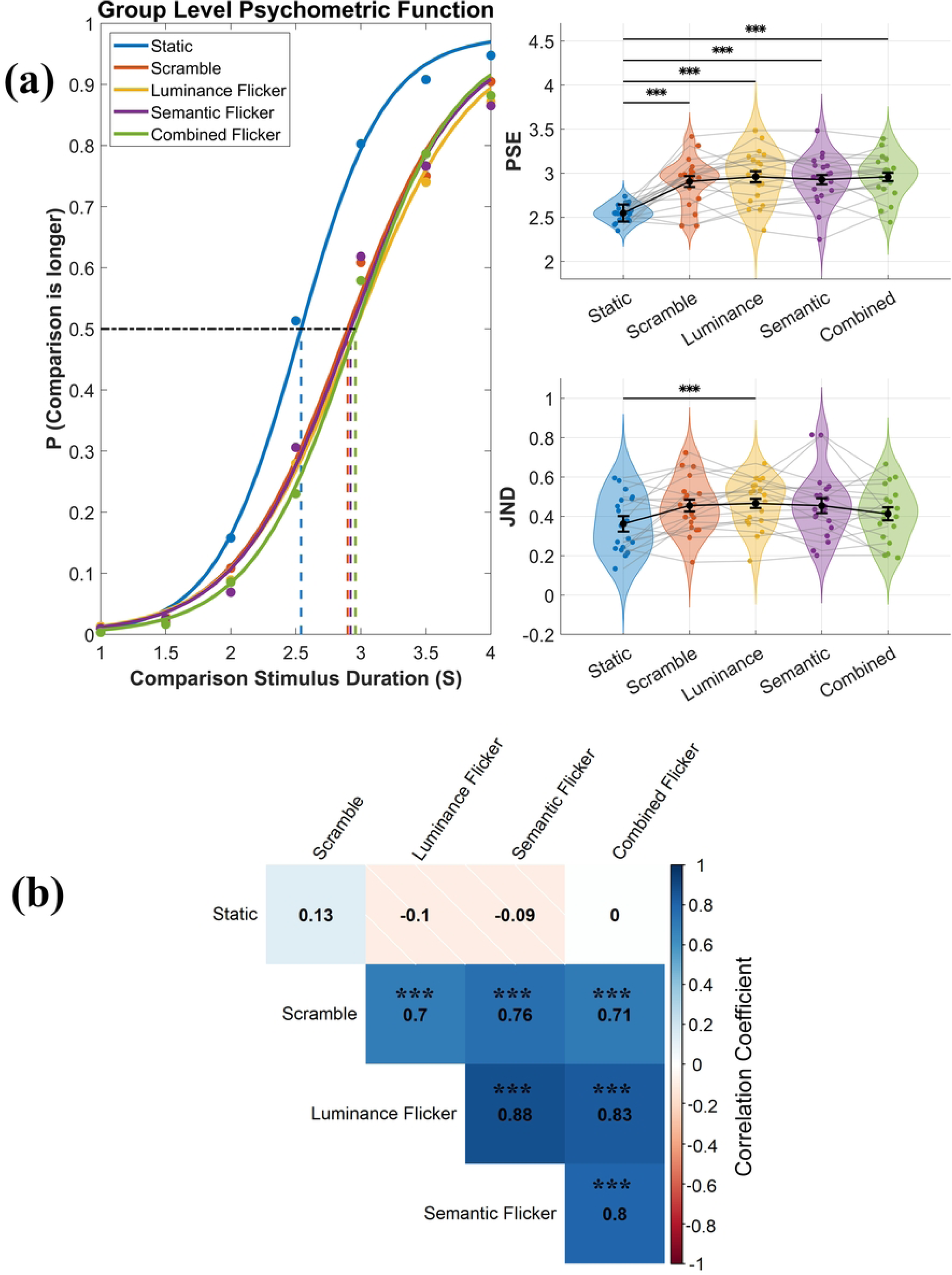
Psychometric function fit. (a) PSEs in the flickering and the scramble conditions shifted significantly to the right relative to the static baseline condition, indicating a time dilation effect. However, no significant difference in PSE was observed either within the flickering conditions or between the flickering and scramble conditions. Analysis of JND indicated the temporal sensitivity was worse in the luminance flicker condition relative to the baseline. (b) represents the PSE correlogram. The correlation pattern among experimental conditions revealed that the time dilation effect was strongly correlated withing flickering conditions and between the flickering and scramble conditions. This points to a shared mechanism driving the time dilation effect which is already present at the level of scramble condition (see text). Error bar indicates 95% confidence interval. *** p < 0.001

To further qualify this finding, we assessed the correlation of PSEs between conditions across subjects. The results are shown in a correlogram in Fig 6b. As is evident, the PSEs in the scramble condition were significantly correlated with the flickering conditions. This means subjects who judged the scrambles to last longer consistently also judged the flickering conditions to last longer. Thus, irrespective of the robust entrainment in flickering conditions, a shared mechanism between flickering and scramble conditions resulted in consistent overestimations across subjects. This shared mechanism is likely related to the scrambling itself, which is a feature present in all other flickering conditions (Fig 3c). This finding highlights the effect of steady activation in V1/V2 or motion-sensitive areas (see Discussion).

Moreover, the multiple linear regression assessing the relation between PSE and the 2-Hz ITPC was insignificant (*F* (9, 85) = 0.37, *p* = 0.94, R^2^ = 0.03). The follow-up single regression analysis revealed that the correlation between PSEs and 2-Hz ITPC values was not significant in any of the conditions (Fig 7). This lack of correlation was also observed between the harmonic frequencies and PSEs (data not shown). Overall, these findings point to a clear separation between behavioral time dilation effect and the robust entrainment observed in the flickering conditions.

**Fig 7.**
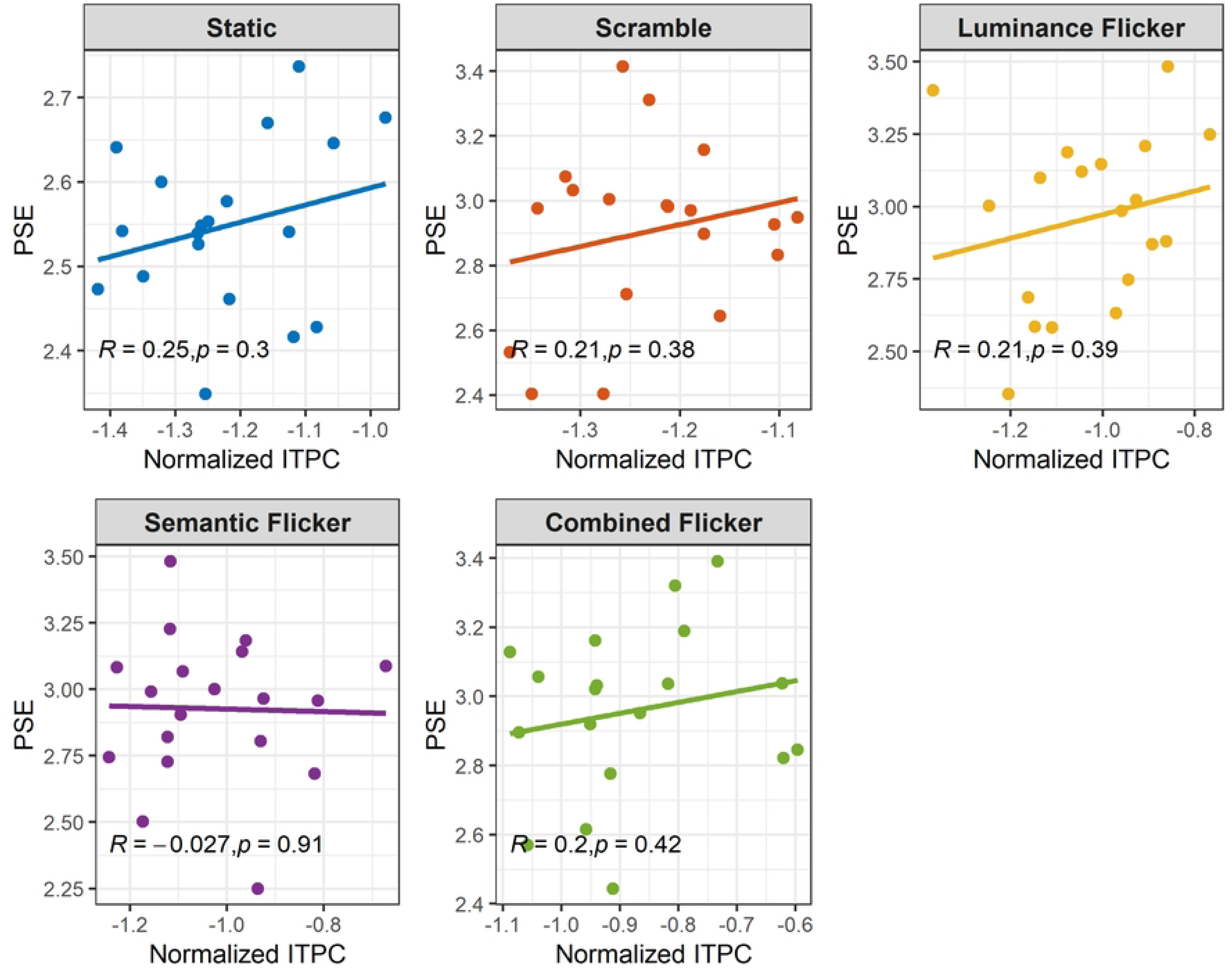
Correlation between PSE and normalized (probit-transformed) 2-Hz ITPC values across subjects. No significant correlation was found in any of the conditions. For correlation between 2-Hz ITPC and JND, see supplementary Fig S2.

Furthermore, the analysis of JND values across conditions revealed that temporal sensitivity was poorer in the luminance flicker condition relative to the static baseline (t (18) = 3.8, *p* = 0.013, BF10 = 29.62). From a frequentist point of view the temporal sensitivity was comparable among experimental conditions (*p* > 0.05). Nevertheless, the Bayse factor revealed that there was moderate evidence for poor temporal sensitivity in scramble (BF10 = 7.38) and semantic flicker (BF10 = 5.88) relative to the baseline static condition. Importantly, there was also moderate evidence suggesting the temporal sensitivity was worse in the luminance flicker relative to the combined flicker condition (BF10 = 5.16, see discussion).

Lastly, the multiple linear regression revealed that the JND and 2-Hz ITPC values were not correlated in any of the conditions (*F* (9, 85) = 0.06, *p* = 1, R^2^ = 0.006; Fig S2). Therefore, these results point to an absence of a relationship between temporal sensitivity and entertainment.

## Discussion

The aim of the current study was to explore whether semantic-based flickers targeting higher-level regions of the ventral stream (IT) can induce notable FITD. Although orthodox luminance-based flickers have previously been used to probe the function of ventral stream (e.g., [42,75]), their interpretation is not straightforward. In such a methodology, since the semantic object representations and the overall luminance/contrast are simultaneously modulated at a base frequency, both higher- and lower-level sections of the visual hierarchy would be concomitantly entrained [33]. Alternatively, SWIFT by preserving the low-level visual properties (global luminance, contrast, and local distribution of spatial frequencies) across the frames, does not entrain the lower-level visual areas [62]. Rather, by rotating the local contours’ orientations in the wavelet domain, it regularly makes the original semantic image recognizable at a fixed frequency [33,62]. Nonetheless, the individual pixels across the frames are equally modulated at a definite frequency spectrum (akin to white noise with a low frequency cut-off, Fig 3a), creating ripple-like motions [33]. To control the potential time distortions rendered by this confounder, we created a condition that included SWIFT scrambles without any recognizable images (Fig 1c, see Method). To compare semantic FITD with the classic FITD, an additional luminance-modulated scramble sequence was created (Fig 1d). Moreover, luminance-modulated semantic flickers were additionally created to probe whether simultaneous entrainment of lower and higher level of the visual hierarchy may additively increase the size of the FITD (Fig 1b). Importantly, in this experimental design on every trial, each of the aforementioned conditions were compared with a static image from which they were derived (Fig 2). This served to ensure that the low-level properties between the standard and comparison stimuli in the duration comparison task were identical. Moreover, static recognizable images were used as comparison stimuli to ensure semantic FITD, if any, is solely resulting from the entrainment of higher-level ventral regions and not from the stimuli’s semantic salience per se.

Our EEG results confirmed that only semantic, luminance, and combined flicker conditions resulted in significant phase clustering. Thus, as expected, the scramble condition did not entrain neural oscillations. The EEG topographic maps (Fig 4a) of luminance flickers align with previous studies using luminance-modulated flickers reporting phase clustering mostly around primary occipital areas with additional contributions from frontal regions (e.g., [36–40]). Moreover, the semantic flickers showed a topography similar to previous studies using SWIFT paradigm [63,70]. The lateralized activation of the right occipitotemporal and temporoparietal electrodes aligns with studies showing category-selective activation in the right temporal regions for faces using SSVEPs [75], face-selective event-related potentials (e.g., N170 [76]; see [77] for a review), and selective right occipitotemporal MEG components for faces and houses [78]. Therefore, the right lateralized phase clustering observed for semantic flickers may result from entrainment of house- and face-selective cortical regions, such as the parahippocampal place and fusiform face areas [62,79]. Lastly, as anticipated, the combined flicker condition resulted in phase clustering across a wide range of topographic areas implying simultaneous engagement of the lower- and higher-level visual regions.

Despite the clear phase clustering observed in luminance, semantic, and combined conditions (Fig 4b & Fig 5), the behavioral data did not show any significant time dilation difference in these conditions relative to the non-entraining scramble condition (Fig 6a). Moreover, the correlation between PSEs and the phase clustering values (at the base frequency and harmonics) across subjects was not significant in any of the conditions (Fig 7). The same insignificant correlations were also obtained between JNDs and phase clustering values in all conditions for the base frequency and its harmonics (Fig S2). These results point to a clear distinction between the observed time dilation and the entrained neural oscillation in luminance, semantic, and combined flicker conditions. Importantly, the significant correlation of subjects’ PSEs between all four conditions (Fig 6b) points to a common mechanism governing time dilation in these conditions which is already present at the level of SWIFT scrambling. This rather unexpected finding implies that the SWIFT scrambling, by steady activation of the lower-level visual [62] and potentially motion sensitive regions, could override the classic FITD effect. This further challenges our current understanding of the mechanisms underlying Flicker- and motion-induced time dilations. Motion-induced time dilation, which is a temporal illusion closely related to FITD, has long been documented [80,81]. However, the controversy surrounding motion-induced time dilation lies in whether the primary driving factor of this illusion is speed or temporal frequency [48,82–85]. In their study, Kanai et al. [48] reduced motion-induced time dilation to FITD showing that the sheer modulation of temporal frequency is the major factor causing dilation, whereas factors such as speed, coherence, and spatial frequencies only have minor intermediary roles. Kaneko & Murakami [82] by closely analyzing the task used by Kanai et al. [48] discovered that the configuration of Kanai et al.’s task had a notable global modulation of luminance, leading the temporal frequency to play a major role in their reported finding. However, our results (Fig 6a) indicate that a global modulation of luminance could not further modulate the time dilation effect which was already engendered by motion and/or steady activation of lower-level visual areas (i.e., scramble condition [62]).

The scramble and the luminance flicker conditions, despite phase clustering differences, primarily engage lower-level visual areas (V1 & V2 [21,41,62]). In the same vein, studies have shown that modulation of lower-level visual areas (V1) with Random Noise Stimulation (tRNS) can dilate the subjective perception of time [9]. Although the frequency distribution of the individual pixel modulation in SWIFT resembles white noise, a previous fMRI study has shown that SWIFT can steadily activate V1 and V2 areas [62]. Thus, whether SWIFT scrambling and tRNS effect on V1 in time dilation are tapping the same mechanism (potentially enhancing cortical excitability [86]) remains to be investigated. On the other hand, owing to the ripple-like motions, ubiquitously present both in the scramble and luminance flicker conditions, it is likely that motion processing areas such as V5/MT [87] have been equally engaged in these conditions. This consequently implies that such higher-level visual areas across the dorsal pathway may have the strongest leverage in causing time dilation effects. Indeed, perceived motion is highly time dilative: even static images with implied motion or illusory jitter can induce time dilation [88,89]. Thus, one may conjecture that flickers dilate time as much as they engage motion sensitive areas. In fact, there is some evidence suggesting luminance-modulated flickers lead to major engagement of V5/MT areas [41]. This compelling speculation suggests that FITD and motion-induced time dilation may be two sides of the same coin, warranting further clarification by future research.

Notwithstanding, the current study was designed to explore whether the entrainment of higher-level ventral visual regions influences FITD. Our results partially rule out this possibility, suggesting that in the presence of steady activation of lower-level visual areas and/or motion sensitive regions, ventral engagement cannot further expand the subjective time. It is important to note that, in our design, each trial involved comparing each of the experimental conditions (static, scramble, luminance, semantic, and combined flickers) to the static image from which their sequence was derived (see Method). This approach was intended to ensure that any time dilation effect observed in the semantic and combined conditions could not be attributed to the *salience* of the semantic images per se; rather, it would purely result from the entrainment of higher-level ventral regions. Had we chosen a different comparison stimulus (e.g., scramble), any time dilation effect in semantic and combined conditions could have been equally attributed to both the salience of the semantic images and the entrainment. Nonetheless, the major limitation of the current study is that we could not find out whether, in the absence of motion, entrainment of ventral regions can induce time dilation. Future development in image processing techniques as well as brain stimulation studies may provide further insights into this issue.

In general, our results are not in agreement with the neural entrainment account of FITD [27,30]. Yet, it is worth noting that, by design, the current study was focused on the supra-second time range. It is suggested that sub- and supra-second ranges rely on distinct timing mechanisms [90–92], and positive correlation between the amplitude of SSVEPs and the magnitude of FITD so far has only been reported for the sub-second ranges [27]. Thus, in line with a previous study [26], our results point to a clear distinction between SSVEPs and FITD in the supra-second range, further constraining the neural entrainment account of FITD.

The results obtained in the current study do not seem to align with the other major account of FITD, which revolves around the concept of subjective salience. To our knowledge, only two studies have assessed the effect of salience on FITD, and each has offered their own interpretation of salience. Herbst et al. [26] considered salience to be a function of human visual system’s flicker sensitivity (peaking at 8-15 Hz [93]). Thus, from their perspective, salience is defined as the degree to which a flicker can consciously be perceived as flickering. On the other hand, by manipulating the number of visible flickers on the screen, Li et al. [28], found robust effect of salience on FITD. Thus, in their study, salience was defined as the interaction between flicker visibility and numerosity. Nonetheless, in the current study, we neither manipulated the range of frequencies, nor the numerosity of flickers. Hence, none of the abovementioned definitions of salience directly applies to the results obtained in our study.

The dilatative effect of salience on time perception seems unanimously agreed upon [10]. However, the concept of salience in time perception studies is rather ill-defined, with different researchers interpreting divergent range of properties as salient (e.g., [44,94–98]). Perhaps the broader theoretical framework of *processing principles* [10] has provided the overarching definition of salience. In this framework, salience is determined by the extent to which the clarity (the information extraction and vividity) of a percept is enhanced by a ‘change’ or its ‘standing out’ from the background context. Based on this definition of salience, in this study, the combined flicker condition should have resulted in greater time dilation than the scramble condition. This is because the combined flicker condition both contained the regular presentation of a semantic image and a regular modulation of luminance. Hence, this in-phase modulation should have resulted in forming a clear representation of this condition in the brain relative to pure scramble (as it is evident by the widespread phase clustering observed for the combined flicker condition, Fig 4 & 5). Therefore, our results do not support the definition of salience provided by the processing principles framework [10]. Moreover, it is worth noting that our results cannot be straightforwardly reconciled with the recently proposed accounts of perceptual content [58,94] and memorability [59] in time perception. If one assumes the combined, or even the semantic and luminance flicker conditions, had higher perceptual content relative to the pure scramble condition, one would expect relatively greater dilation for these conditions. Similarly, it is unclear why the scramble condition would have the same memorability as the semantic flicker condition, resulting in comparable time dilations [59].

One possibility is that the modulation of the individual pixels (Fig 3a) and/or the ripple-like motions [33] as the by-product of SWIFT scrambling had saturated the time dilation effect. However, this suggestion seems unlikely. This is because if saturation were present, the temporal sensitivity (i.e., the discrimination threshold as measured by JND) should have remained the same in saturated conditions relative to the baseline. Yet, as shown in Fig 6a, the JND, specifically, for the luminance condition was significantly worse than the baseline static condition (similar to findings reported by [99]). Moreover, moderate evidence existed for the difference of JND between luminance and combined flicker condition (BF10 = 5.16). Nevertheless, to formally investigate whether saturation has smeared our results, we conducted a supplementary analysis. For each subject, we computed the standard deviation of PSEs across the four conditions of scramble, luminance, semantic, and combined flickers. We then set a moderate threshold of 0.1 seconds and subjects whose standard deviation fell below this value were removed. This means for the remaining subjects (12 out of 19), 68% of PSEs in the abovementioned conditions fell within at least ±0.1-seconds range around each subject’s mean. Saturation, if any, thus was controlled for these remaining subjects who showed notable within-subject variance across the scramble and flickering conditions. Repeating the same analysis, conducted on the entire sample, for this subset of subjects did not alter any patterns of significance. Importantly, the Bayse factors indicating null differences across experimental conditions were comparable to those obtained from the analysis of the whole sample (see supplementary materials).

Lastly, we conducted a supplementary event-related potential (ERP) analysis to assess whether the contingent negative variation (CNV) and the N1/P2 amplitudes were modulated differently across conditions (Fig S5). These ERP components were chosen based on their implied role in time perception studies [100,101]. The ERPs across the conditions were nearly identical (see supplementary materials). Therefore, neither the CNV nor the N1/P2 amplitude was significantly different among conditions. This finding is in line with the null results reported for CNV amplitude differences between flickering and static conditions [100]. Inevitably, the correlations of these ERP components with PSEs were as well insignificant in all conditions. Additionally, we conducted an exploratory time-frequency power analysis (Fig S6). In the flickering conditions, a broadband desynchronization of alpha and beta oscillations was observed, which was absent in the scramble condition. Nevertheless, relative to the static baseline condition, this broadband desynchronization did not reach statistical significance (p > 0.05, see supplementary materials).

## Conclusion

In this pre-registered study, we aimed to explore whether the higher-level ventral visual regions contribute to FITD using SWIFT technique. However, we found that the SWIFT scrambling per se resulted in a time dilation effect which was not further amplified by periodic luminance or semantic modulations. This rather surprising finding may suggest that FITD, and motion-induced time dilation may rely on the same mechanism based on the activation of areas such as V5/MT. On the other hand, our results indicate that, at least at the presence of steadily activated lower-level visual and/or motion sensitive areas, periodic activation of ventral visual regions does not contribute to FITD. Brain stimulation techniques such as transcranial magnetic stimulation (TMS) and tRNS have provided noteworthy findings regarding the role of primary visual (V1) and V5/MT motion areas in time perception [4,9,102,103]. Studying FITD using brain stimulation techniques, in conjunction with results obtained from image processing methods (such as SWIFT), would enhance our understanding of how and which levels of the visual hierarchy underlie FITD. Lastly, while our results do not align with the neural entrainment account of FITD, they also do not support the processing principles definition of salience. Systematic research is needed to clarify what salience in time perception is and how one can objectively and operationally define and measure it.

## Data availability

Reported data and analysis scripts from all experiments are openly available on the Open Science Framework (https://osf.io/tj3b7/).

